# Beyond Phase Estimation: A Multidimensional Gating Framework for Robust Real-Time Closed-Loop Neural Stimulation

**DOI:** 10.64898/2026.05.26.727783

**Authors:** Wenjia Zheng, Lu Shen, Biao Han

## Abstract

Neural oscillatory phase is widely used as a control variable in real-time closed-loop stimulation, yet its validity under strict causal constraints and noisy conditions has rarely been systematically examined. We introduce a Multidimensional Gating Framework (MGF), a plug-in and estimator-agnostic module that determines whether phase information should be admitted into control by evaluating instantaneous amplitude, narrowband signal-to-noise ratio (SNR), and spectral peak ratio (PR) within a strictly causal window. Using causal streaming replay on a public resting-state EEG dataset, we benchmarked Hilbert based phase estimation and endpoint-corrected Hilbert estimation with and without MGF. Among feasible subjects, MGF significantly reduced phase dispersion for both estimators, while robustly suppressing catastrophic phase errors. In contrast, ungated approaches exhibited systematic failures under the same conditions.

## Introduction

Neural oscillatory phase has long been considered a key variable for characterizing fluctuations in perception and behavior, particularly in the alpha band (approximately 8–12 Hz). A substantial body of work has shown that, even when stimulus intensity is held constant, prestimulus alpha phase predicts visual awareness, perceptual thresholds, and response probability (Ai & Ro, 2014; Busch et al., 2009; Riecke et al., 2015). These findings have motivated theories of perceptual rhythmicity, which propose that cognitive processing is modulated by the phase structure of endogenous neural oscillations.

Within this framework, phase-locked closed-loop stimulation has emerged as a powerful approach for testing the causal role of oscillatory phase. Closed-loop systems aim to track neural activity in real time and trigger stimulation contingent on the estimated phase, thereby elevating phase from a correlational descriptor to an explicitly manipulable control variable. With advances in hardware and real-time signal processing, such approaches have been increasingly applied across perceptual, cognitive, and clinical domains (Harlow et al., 2024; Hebron et al., 2024; Ngo & Staresina, 2022; Schreglmann et al., 2021). If oscillatory phase indeed plays a causal role, stimulation delivered at different phases should yield systematic and predictable differences in outcomes.

However, most existing closed-loop studies rely on an implicit but critical assumption: that phase is always usable as a control variable as long as it can be numerically estimated. This assumption is typically justified in offline analyses, where phase estimation can exploit zero-phase filtering, full temporal context, and post hoc trial selection, yielding stable and internally consistent phase definitions. In contrast, real-time closed-loop systems must operate under strict causal constraints, with phase estimates depending only on past data and being subject to transmission delays, finite analysis windows, and measurement noise.

To mitigate endpoint distortions under causal constraints, the endpoint-corrected Hilbert transform (ecHT) was proposed to improve numerical stability in online phase estimation (Schreglmann et al., 2021). While ecHT addresses estimator-level numerical bias, it does not resolve a more fundamental methodological question: whether phase should be used at all when the underlying oscillation is weak, transient, or strongly non-stationary. Indeed, multiple studies have failed to observe robust phase effects (Benwell et al., 2017; Melcón et al., 2024; Michail et al., 2022; Ruzzoli et al., 2019), and systematic analyses show that real-time phase detection errors depend strongly on spectral structure and signal quality, deteriorating substantially under non-ideal conditions (Liufu et al., 2025).

Spontaneous EEG oscillations are typically low in amplitude, intermittent, and highly non-stationary. Under such conditions, an algorithm may output a numerically well-defined phase angle that does not correspond to a methodologically interpretable oscillatory state. If a closed-loop system continues to execute phase-locked triggering during periods of weak or spectrally ambiguous oscillations, triggered events will mix samples drawn from heterogeneous neural states. This mixture degrades phase alignment reliability and undermines the interpretability of phase-locked stimulation as a causal intervention. Consequently, the central challenge of closed-loop phase stimulation is not merely to improve phase estimation accuracy, but to determine when phase should not be used. Although some real-time studies have imposed amplitude thresholds at trigger time (Vigué-Guix et al., 2022), the methodological rationale for such practices has not been formally articulated. Here, we explicitly treat phase usability as a prerequisite for real-time closed-loop control and introduce the Multidimensional Gating Framework (MGF). Before each candidate trigger, MGF evaluates multiple indicators of oscillatory salience within a strictly causal window, including analytic amplitude, narrowband signal-to-noise ratio, and spectral peak ratio. Phase information is admitted into the triggering logic only when the current oscillatory state satisfies explicit interpretability criteria. Importantly, MGF functions as an estimator-agnostic external wrapper: it leaves the internal implementation of phase estimators unchanged while imposing a general-purpose reliability constraint on the closed-loop pipeline.

Using a publicly available resting-state EEG dataset (Hatlestad-Hall et al., 2022), we implemented a causal streaming replay framework that faithfully reproduces real-time data transmission and latency constraints. Within this framework, we systematically evaluated four closed-loop phase-triggering strategies, combining two phase estimators (Hilbert and ecHT) with and without MGF. This 2 × 2 design allows us to test whether MGF exerts consistent effects across estimators, rather than comparing the estimators themselves.

Our results show that explicitly modeling phase usability is a necessary prerequisite for effective real-time closed-loop phase stimulation. By restricting phase-based control to states with sufficient oscillatory salience, MGF enhances trigger reliability and mitigates catastrophic phase mismatches. This framework thus delineates a clear methodological boundary for treating oscillatory phase as a causal control variable. The contributions are as follows:

1. We demonstrate that under strict real-time causal constraints, the assumption that “phase is always usable” is untenable. We reformulate the core problem of closed-loop phase stimulation from optimizing phase estimation accuracy to determining the conditions under which phase can serve as a valid control variable, explicitly distinguishing numerical stability from methodological interpretability.
2. We introduce the Multidimensional Gating Framework (MGF) as an estimator-agnostic external wrapper that evaluates multidimensional oscillatory salience within a strictly causal window and explicitly determines whether phase should enter the control loop. This design provides an essential system-level reliability constraint for real-time closed-loop phase stimulation.
3. Using causal streaming replay and systematic comparisons across two phase estimators (Hilbert and ecHT) with and without MGF, we show that MGF robustly improves trigger reliability in spontaneous EEG and substantially reduces the long-tail risk of catastrophic phase mismatches, establishing its role as a necessary component of real-time closed-loop phase stimulation.

## Method

### Overview

We propose and validate a Multidimensional Gating Framework (MGF) for real-time closed-loop phase-locked stimulation. MGF is designed as an estimator-agnostic wrapper that operates upstream of phase estimation, explicitly determining whether instantaneous phase is methodologically interpretable under strict causal constraints. As illustrated in Figure 1, the framework evaluates multiple online measures of oscillatory salience and only admits phase information into the triggering logic when all necessary conditions are met. This design elevates the question of when phase should not be used from an implicit assumption to a core component of closed-loop control.

**Figure 1:**
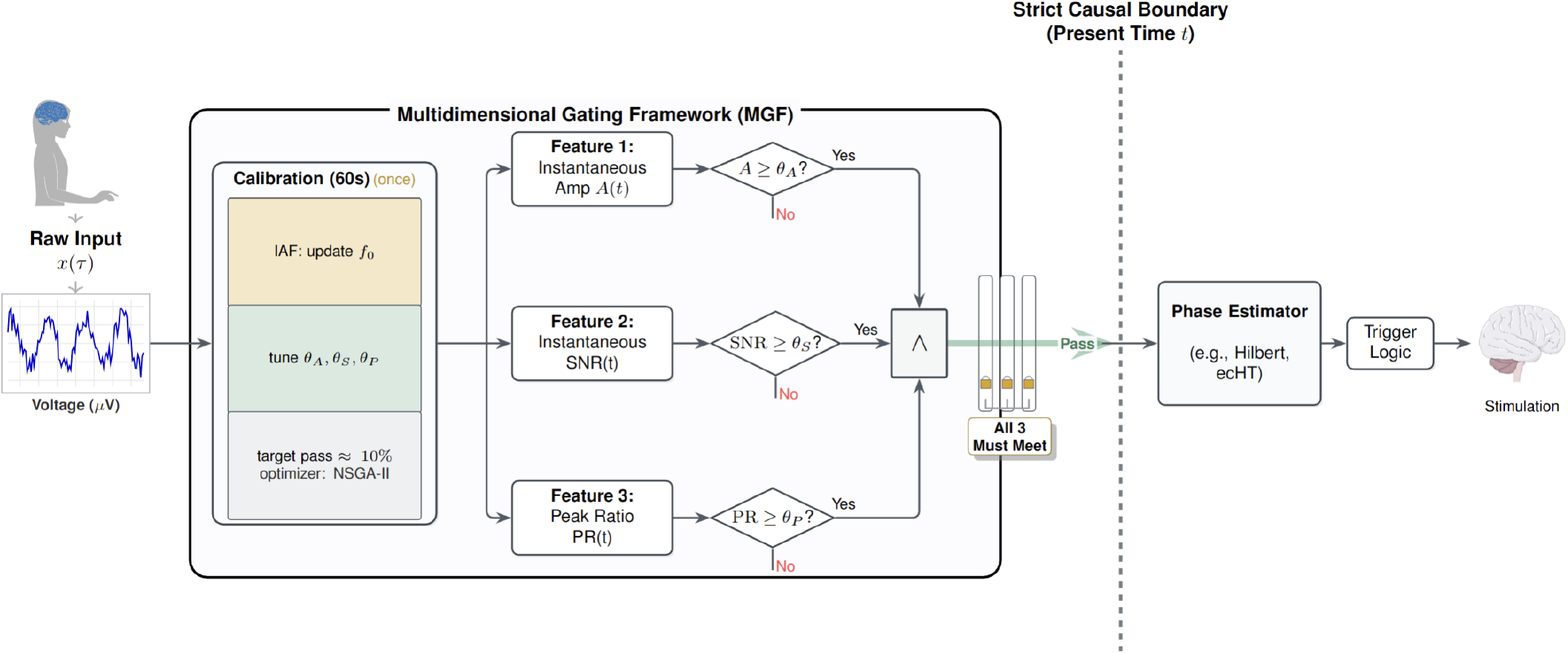
Overview of the Multidimensional Gating Framework (MGF). Raw EEG input *x* (*τ*) is processed under strict causal constraints. After a one-time calibration that sets subject-specific thresholds with a target pass rate of *q* = 10%, three online features are jointly evaluated: instantaneous amplitude *A* (*t*), signal-to-noise ratio (SNR), and peak ratio (PR). Phase estimation and stimulation are enabled only when all criteria are satisfied. The dashed vertical line indicates the strict causal boundary at time *t*.

### Problem Formulation and Theoretical Limits

#### Strictly causal formulation

Let the observed signal be

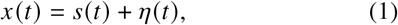

where *s* (*t*) denotes a target narrowband oscillatory component that may be transient and nonstationary, and *η* (*t*) represents broadband background activity and noise. In real-time closedloop operation, the system at time *t* has access only to past samples *x* _(− ∞,*t*)_. Accordingly, an online phase estimator can be written as

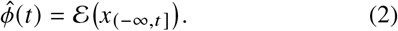

In the closed-loop context, phase is not a descriptive statistic but a control variable used to trigger stimulation. System reliability is therefore determined by the distribution of the phase error

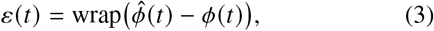

rather than by point-estimation accuracy alone. If this error distribution exhibits heavy tails or multimodality, the system will produce catastrophic phase mismatches with non-negligible probability, even when average performance appears acceptable.

#### Amplitude- and spectrum-dependent limits of phase usability

After strictly causal bandpass filtering, the narrowband signal *y*(*t*) admits the analytic representation

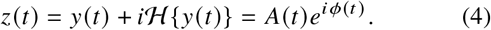

A first-order perturbation analysis shows that phase sensitivity to noise scales inversely with instantaneous amplitude,

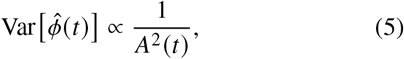

implying that phase estimation becomes ill-conditioned as *A* (*t*) decreases. Thus, during low-amplitude segments, a numerically defined phase may exist but cannot serve as a stable control variable.

Numerical stability alone, however, does not guarantee interpretability. Within a strictly causal window *W*_*t*_ = [*t* −*T, t*], we characterize local spectral structure using the amplitude spectrum |*X* (*f* ; *t*)| and define two online metrics: SNR, the log-ratio of mean alpha-band amplitude to adjacent-band amplitude (in dB), and PR, the ratio of peak to median amplitude within the alpha band. When SNR or PR is low, the target band does not form a unique dominant component, and the observed signal admits multiple plausible narrowband decompositions.

In this regime, compressing the system state into a single phase variable is not identifiable, leading to multimodal and heavy-tailed phase error distributions.

#### Formal link to tail risk

The combined effects of low amplitude and poor spectral identifiability act directly on the error distribution *p* (*ε*). Under these conditions, the probability mass in the tails necessarily increases, yielding the following implication:

#### Proposition (Lower bound on tail risk)

When *A* (*t*) is small or when SNR/PR is low, the conditional distribution *p* (*ε*(*t*) | *x*_(−∞,*t* ]_) exhibits increased tail mass, and the lower bound of

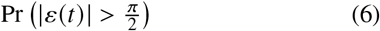

is elevated.

This result implies that catastrophic phase mismatches are unavoidable if phase is used indiscriminately during such segments. Consequently, evaluating closed-loop phase stimulation requires not only average accuracy or dispersion metrics, but also explicit quantification of tail risk. This directly motivates the Tail120 (|*ε*| > 120^°^) metric used in the Results.

#### Consequence: explicit modeling of phase usability

Together, these considerations show that phase is not always a valid control variable. Phase usability depends on three online-verifiable necessary conditions:

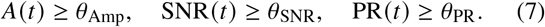

We therefore define a gating variable

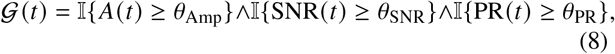

which is evaluated prior to phase estimation. Only when 𝒢 (*t*) = 1 is phase estimation enabled and allowed to enter the triggering logic; otherwise the system explicitly issues a Reject decision. This design constitutes the Multidimensional Gating Framework (MGF).

### Calibration and Parameter Determination

The gating logic defined relies on subject-specific thresholds *θ*_Amp_, *θ*_SNR_, and *θ*_PR_. To determine the optimal parameters, we implemented an adaptive calibration framework based on the Non-dominated Sorting Genetic Algorithm II (NSGA-II). Rather than relying on fixed heuristics, this scheme treats the target joint pass rate (*q* = 0.1) as an explicit constraint and performs a heuristic search over the multidimensional threshold space to identify subject-specific gating parameters. The optimization balances achieving the desired stimulation throughput with minimizing deviations from conservative baseline threshold values. This procedure enables the MGF system to adapt to inter-subject variability in signal-to-noise characteristics while selectively targeting data segments with reliably high oscillatory salience for each individual.

We define the empirical pass rate under thresholds *θ* = [*θ*_Amp_, *θ*_SNR_, *θ*_PR_]^⊤^ as

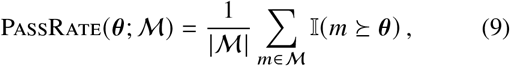

where ⪰ denotes component-wise satisfaction of the gating criteria. This quantitative formulation serves as the objective function for the subject-specific gating calibration described in Algorithm 1.

#### Algorithm 1

NSGA-II Calibration of Subject-Specific Gating Thresholds

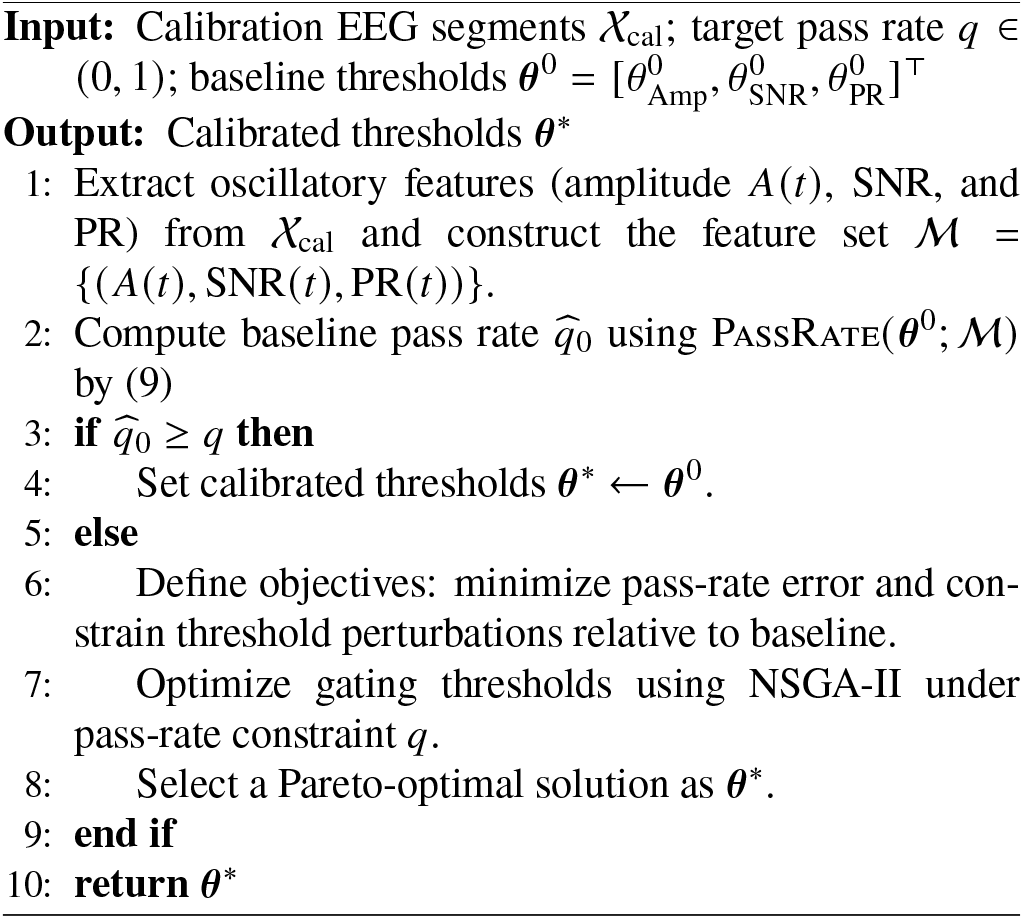

### Algorithmic Instantiation and Causal Simulation

We instantiated the proposed Multidimensional Gating Framework (MGF) as an estimator-agnostic control layer that explicitly regulates whether instantaneous phase information is allowed to enter the closed-loop triggering logic. MGF operates upstream of phase estimation and can be coupled to arbitrary causal phase estimators without modifying their internal structure.

To demonstrate this estimator-agnostic property, MGF was instantiated with two representative phase estimation backends: the standard causal Hilbert transform and the endpoint-corrected Hilbert transform (ecHT). For each estimator, performance was evaluated with and without the proposed gating mechanism, yielding four experimental conditions used solely for comparative evaluation. Importantly, the underlying phase estimators were identical across gated and ungated conditions; any performance differences therefore reflect the effect of MGF rather than estimator-specific modifications.

As a prerequisite for online deployment, gating thresholds were determined during a brief calibration phase prior to online operation as shown in Algorithm 1. When predefined baseline thresholds did not meet the target pass rate, they were adaptively refined using a multi-objective genetic algorithm NSGA-II (Deb et al., 2002). This calibration step was performed once and did not alter the estimator-agnostic structure or the real-time triggering logic of MGF.

To evaluate performance under realistic conditions, we implemented a streaming replay that strictly reproduces real-time causality and latency constraints on offline data. Trigger decisions were made online using only past samples, while reference phases were computed offline solely for evaluation. Specifically, the ground-truth phase was obtained by applying zero-phase bandpass filtering followed by Hilbert transform to the entire recorded session, ensuring that reference values were free from causal delays and endpoint artifacts.

### Evaluation Metrics and Statistical Analysis

#### Data Aggregation and Metrics

Analysis was performed at the subject level to ensure statistical independence. For each subject and configuration (MGF vs. NoMGF), we computed three performance metrics: the hit rate, defined as the proportion of triggers that resulted in valid stimulation; the circular standard deviation (CircStd), a measure of phase precision, computed only for subjects with sufficient trigger counts (*N*_trig_ ≥ 5) to ensure numerical stability; and Tail120, defined as the proportion of stimulation events with absolute phase error exceeding 120^°^.

#### Statistical Framework

All performance metrics (Hit Rate, CircStd, and Tail120) were first computed at the stimulation-event level and subsequently aggregated within subjects for each experimental condition. These subject-level metrics constituted the dependent variables for all subsequent statistical analyses; no metric was computed within the statistical models themselves.

To validate the effects of the proposed Multidimensional Gating Framework (MGF), we employed a two-tier statistical analysis strategy combining linear mixed-effects modeling (LMM) with robust non-parametric paired tests.

#### Linear Mixed-Effects Analysis

Linear mixed-effects models were used to quantify the systematic interaction between the proposed gating strategy and the underlying phase estimation algorithm. For each subject-level metric, we fitted the following model:

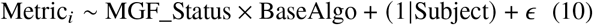

where MGF_Status denotes whether MGF was applied (MGF vs. NoMGF), and BaseAlgo denotes the base phase estimator (ecHT vs. Hilbert). This factorial design explicitly tests for an interaction effect, indicating whether the benefit of MGF depends on the choice of phase estimator. A random intercept for each subject was included to account for within-subject correlations across conditions. Statistical significance was assessed using Satterthwaite’s approximation for degrees of freedom.

#### Robust Paired Comparisons

As a complementary validation, paired Wilcoxon signed-rank tests were conducted to directly compare MGF against NoMGF within each estimator family (e.g., MGF(ecHT) vs. NoMGF(ecHT)). These analyses provide a distribution-free assessment of within-subject differences and serve as a robustness check against potential deviations from parametric assumptions.

All tests were two-sided, and *p*-values were adjusted for multiple comparisons using the Holm–Bonferroni procedure. Effect sizes were quantified using the rank-biserial correlation (*r*_*rb*_), with conventional thresholds for interpretation. Confidence intervals (95%) for median differences were estimated using the Hodges–Lehmann estimator.

## Results

### Performance Overview

All analyses were based on data from *N* = 86 subjects who satisfied the criteria for data completeness and a minimum trigger count (*N*_trig_ ≥ 5). The original dataset comprised *N* = 110 subjects; the reduction in sample size reflects statistical constraints inherent to within-subject paired analyses, which require stable and estimable metrics across all comparison conditions, rather than any post-hoc selection based on performance or statistical significance.

The Multidimensional Gating Framework (MGF) exhibited distinct performance characteristics depending on the underlying phase estimator. Descriptive and inferential statistics for all primary metrics are summarized in Table 1.

**Table 1:**
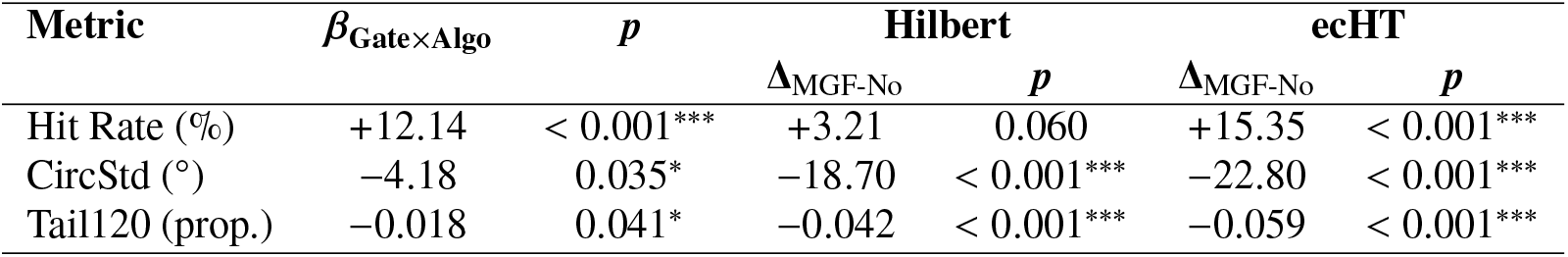
Fixed effects and simple effects from linear mixed-effects models (*N* = 86). Interaction terms correspond to MGF × Algorithm. Simple effects report estimated differences between MGF and NoMGF within each estimator. Significance levels: ^∗^ *p* < 0.05, ^∗∗∗^ *p* < 0.001.

| Metric | $\beta_{\text{Gate} \times \text{Algo}}$ | $p$ | Hilbert | | ecHT | |
| --- | --- | --- | --- | --- | --- | --- |
| | | | $\Delta_{\text{MGF-No}}$ | $p$ | $\Delta_{\text{MGF-No}}$ | $p$ |
| Hit Rate (%) | +12.14 | < 0.001*** | +3.21 | 0.060 | +15.35 | < 0.001*** |
| CircStd ($^\circ$) | -4.18 | 0.035* | -18.70 | < 0.001*** | -22.80 | < 0.001*** |
| Tail120 (prop.) | -0.018 | 0.041* | -0.042 | < 0.001*** | -0.059 | < 0.001*** |

### Interaction Effects Between Gating and Estimator

To quantify the effects of gating and base phase estimator, we fitted linear mixed-effects models (LMMs) separately for Hit Rate, circular standard deviation (CircStd), and Tail120. Each model included fixed effects of Gating (MGF vs. NoMGF), Base Algorithm (ecHT vs. Hilbert), and their interaction, with a random intercept for subject. Models were fitted using restricted maximum likelihood (REML), and inference relied on Satterthwaite-approximated degrees of freedom.

Across all three metrics, significant Gating × Algorithm interactions were observed, indicating that the effect of MGF was not uniform across estimators but depended systematically on the properties of the underlying phase estimator.

#### Hit Rate

For Hit Rate, the LMM revealed a significant interaction between Gating and Algorithm (*β* = 12.14, *t* = 5.06, *p* < 0.001). Simple-effects analysis showed that, for the standard Hilbert estimator, MGF produced a modest numerical increase in Hit Rate that did not reach significance in the parametric model (estimated Δ = +3.21 percentage points, *p* = 0.060). In contrast, when applied to ecHT, MGF yielded a large and highly significant improvement in throughput (estimated Δ = +15.35 percentage points, *p* < 0.001). This pattern indicates that while MGF enforces signal-quality constraints, substantial gains in valid stimulation throughput emerge primarily when combined with a predictive phase estimator.

#### Phase Precision (CircStd)

For phase precision, CircStd exhibited a strong main effect of gating as well as a significant interaction (*β* = −4.18, *t* = −2.12, *p* = 0.035). MGF significantly reduced angular dispersion for both estimators (main effect *p* < 0.001), but the magnitude of this reduction differed across algorithms.

Specifically, the estimated marginal mean of CircStd was reduced by approximately 18.7° for Hilbert and by 22.8° for ecHT, indicating a larger precision gain for the predictive estimator.

#### Tail Risk (Tail120)

Analysis of catastrophic phase errors, quantified by Tail120, also revealed a significant Gating × Algorithm interaction (*β* = −0.018, *t* = −2.06, *p* = 0.041).

MGF effectively suppressed heavy-tailed phase errors for both estimators, but the reduction in the proportion of large errors was significantly greater when gating was combined with ecHT than with Hilbert.

### Robust Paired Validation and Bias Analysis

To validate the LMM results without parametric assumptions, we conducted paired Wilcoxon signed-rank tests within each estimator family, using two-sided tests with Holm correction across metrics. In addition, phase bias was analyzed exclusively in this non-parametric framework due to its sign dependence and asymmetric distribution.

#### Hilbert Estimator

For the Hilbert estimator, MGF significantly improved phase precision (median CircStd Δ = 16.9°, *p* < 0.001, *r*_*rb*_ = −0.93) and reduced tail risk (median Tail120 Δ = −0.05, *p* < 0.001). The non-parametric analysis also detected a small but significant increase in Hit Rate (median Δ = +1.3%, *p* = 0.023, *r*_*rb*_ = 0.28), consistent with a modest throughput gain not reaching significance in the parametric LMM. Notably, MGF induced a substantial negative shift in phase bias for Hilbert (median Δ = −19.6°, *p* < 0.001, large effect), indicating that gating may preferentially select segments associated with systematic phase offsets in this estimator.

#### ecHT Estimator

When applied to ecHT, MGF produced large and consistent improvements across all primary metrics. Hit Rate increased markedly (median Δ = +18.0%, *p* < 0.001, *r*_*rb*_ = 0.96), phase dispersion was strongly reduced (median CircStd Δ = −24.1°, *p* < 0.001, *r*_*rb*_ ≈ −1.00), and Tail120 decreased to near-zero levels. In contrast to Hilbert, phase bias under ecHT was not significantly affected by gating (median Δ = −3.6°, *p* = 0.938), indicating that the predictive estimator maintained calibration stability even under stringent gating constraints.

## Discussion

This study challenges a prevalent yet rarely formalized assumption in the field of real-time closed-loop phase-locked stimulation: that numerically estimated oscillatory phase inherently constitutes an effective control variable. Our results demonstrate that under strict causal constraints, this assumption does not hold. Instead, the usability of phase critically depends on the transient significance and stability of the underlying oscillation. By explicitly modeling this dependency, the Multidimensional Gating Framework (MGF) establishes a fundamental criterion for distinguishing between interpretable and non-interpretable phase control methods.

The core contribution of this work lies in reframing the closed-loop phase stimulation problem from one of estimator accuracy to one of control effectiveness. Previous research primarily focused on enhancing online phase estimation accuracy through algorithmic optimization. While these advances are indispensable, our findings reveal that precise estimation alone remains insufficient. When oscillatory signals are weak, transient, or spectrally ambiguous, even numerically stable estimators may produce phase values lacking causal interpretability. Under such conditions, phase errors exhibit heavy-tailed distributions, leading to rare yet severe failures. MGF directly addresses this failure mode by precisely suppressing phase-based triggering mechanisms when oscillatory conditions violate interpretability criteria.

MGF is implemented as an estimator-agnostic wrapper rather than an algorithm-specific modification. The gating mechanism consistently improves phase accuracy and reduces long-tail error risks across both standard Hilbert transform and endpoint-corrected Hilbert estimators (ecHT). Crucially, the interaction between the gating mechanism and estimator type demonstrates that MGF does not function as a uniform filter. It selectively enables predictive estimators to operate under oscillatory states, ensuring their advantages are reliably leveraged. This design allows MGF to seamlessly integrate with existing and future phase estimation methods without altering their internal architecture.

A key insight from MGF lies in the interpretation of reduced feasibility under gating. Applying MGF inherently excludes time periods or subjects where oscillatory conditions are unsuitable for phase control. This exclusion does not constitute data loss but rather explicitly acknowledges that phase is not always a control-relevant variable. From a causal perspective, excluding invalid trials is preferable to implicit inclusion—the latter confuses heterogeneous neural states and undermines interpretability. Thus, reduced feasibility represents an expected, information-rich outcome of principle-based gating mechanisms.

Tail risk analysis provides compelling corroboration for this interpretation. Average performance metrics may mask rare yet catastrophic errors, particularly challenging in closed-loop stimulation. Our results demonstrate that MGF significantly suppresses extreme phase errors, indicating that the gating mechanism reshapes the error distribution itself rather than merely improving its central tendency. This finding aligns with theoretical predictions—reduced oscillatory significance raises the lower bound of catastrophic phase error probabilities, underscoring the importance of explicitly modeling tail behavior in closed-loop systems.

Several limitations of this study warrant attention: The assessment focused on resting-state EEG, a challenging yet ecologically relevant scenario for phase control. Task-induced or strongly synchronized oscillations may exhibit different feasibility characteristics, though the principle of phase availability remains universal. Additionally, though thresholds were calibrated per-subject, the target pass rate *q* was fixed; fully adaptive or performance-driven thresholding may further enhance robustness. Finally, while causal stream-of-replay simulations enabled real-time operation, future work should explore fully online implementations incorporating hardware-specific constraints.

More broadly, these findings transcend the context of alpha-band EEG stimulation. Any closed-loop system employing estimated latent variables as control signals must address the question of when such variables become well-defined. By transforming rejection into explicit, principled outputs, MGF provides a universal template for real-time neural control systems with integrated methodological safeguards.

In summary, reliable closed-loop phase stimulation requires not only precise phase estimation, but also explicit modeling of phase availability. By mandating oscillatory significance as a control prerequisite, MGF systematically enhances reliability and suppresses catastrophic failures, establishing phase gating as an indispensable methodological component for real-time closed-loop phase stimulation.

## Conclusion

This study demonstrates that reliable real-time closed-loop phase stimulation requires more than accurate phase estimation: it requires explicit modeling of when phase is a valid control variable. By introducing the Multidimensional Gating Framework (MGF), we show that enforcing oscillatory salience as a prerequisite for phase-based triggering systematically improves precision, increases effective throughput, and suppresses catastrophic long-tail errors under strict causal constraints. Importantly, MGF operates as an estimator-agnostic control layer, complementing rather than replacing existing phase estimation methods. More broadly, this work reframes closed-loop phase stimulation as a problem of control validity rather than one of estimation accuracy.

## Acknowledgments

This work was supported by the National Key Research and Development Program of China (2025YFE0213500); the National Natural Science Foundation of China (32471144); the Research Center for Brain Cognition and Human Development, Guangdong, China (2024B0303390003); the MOE Project of Key Research Institute of Humanities and Social Sciences in Universities (22JJD190006); and the Striving for the First-Class, Improving Weak Links and Highlighting Features (SIH) Key Discipline Program for Psychology at South China Normal University.

## Declaration of generative AI and AI-assisted technologies in the manuscript preparation process

An AI-assisted language model (ChatGPT, OpenAI) was used to support English language polishing. All scientific content and conclusions remain the sole responsibility of the authors.

